# Microarray, validation and gene-enrichment approach for assessing differentially expressed circulating miRNAs in obese and lean heart failure patients

**DOI:** 10.1101/2025.02.15.638475

**Authors:** Douglas dos Santos Soares, Amanda Lopes, Mariana Recamonde-Mendoza, Rodrigo Haas Bueno, Raquel Calloni, Nadine Clausell, Santiago Alonso Tobar Leitão, Andreia Biolo

## Abstract

**Background:** Obesity is a risk factor associated with cardiovascular diseases that may lead to heart failure (HF). However, in HF, overweight and obese patients have longer survival than underweight patients, a phenomenon known as the obesity paradox. MiRNAs play a fundamental role in gene regulation and transcription factors involved in obesity and HF. The main objective of this study was to identify and validate differentially expressed circulating miRNAs in HF-obese and HF-lean patients.

**Materials and Methods:** This case-control study was carried out in two parts; the discovery and validation phase. In the discovery phase, plasma samples from 20 HF patients and from 10 healthy controls for microarray analysis were selected. The miRNAs were extracted and then analyzed on the miRNA 4.0 Affymetrix GeneChip array following the manufacturer’s instructions. Raw data were normalized using Robust Microarray Average (RMA), batch effects were adjusted with Surrogate Variable Analysis (SVA), and differential expression analysis was performed with the Limma R package. Differentially expressed miRNAs were ranked based on effect size, p-value, and biological plausibility, and selected for validation. In the validation phase, plasma miRNAs from 80 patients and controls were extracted and miRNAs -451a, - 22-3p, and -548ac were validated by quantitative reverse transcription-PCR. Then, they were subjected to target analysis using miRTarBase release 8.0 and TarBase v8.0. We performed functional enrichment analysis of miRNA targets using pathway annotation from the KEGG Pathway Database using the ClueGO software.

**Results:** MiRNAs -451a, -22-3p, and -548ac were up-regulated in HF-lean and HF-obese groups compared to control, showing an association of these miRNAs with HF. Enrichment analysis showed an association of validated miRNAs with *AKT1*, *MAPK1*, *GRB2*, and *IGF1R* genes. These genes are associated with different biological processes, such as apoptosis, carbohydrate metabolism, neurogenesis, sugar transport, translation regulation, transport, cell cycle, actin cytoskeleton reorganization, cardiac neural crest cell development involved in heart development, and aging.

**Conclusion:** miR-451a, miR-22-3p, and miR-548ac are up-regulated in HF, independent of obesity, and are associated with metabolic, morphological, and functional outcomes. These results might help in the selection of targets for studies aiming to underscore the mechanisms of HF.

## Introduction

Obesity is one of the main risk factors associated with cardiovascular diseases that culminate in heart failure (HF) [1–3]. The higher the body mass index, the greater the risk of developing HF [3]. However, once HF is established, overweight and obese patients have more prolonged survival than patients with low weight and severe obesity [4–7]. This paradox has drawn the scientific community’s attention to better understand the possible mechanisms involved in this scenario. MiRNAs play a key role in gene regulation and there are transcription factors involved in obesity [8] and HF [9]. We performed a case-control study and observed that in HF patients, the presence of obesity is associated with a differential expression of selected miRs and the miR-221/-130b ratio had significant correlations with adiposity parameters [10]. In addition, computational target prediction analysis identified several interrelated pathways targeted by miR-130b and miR-221 with a known relationship with endocrine and cardiovascular diseases [10]. Studying the mechanisms involved in this scenario is of great importance to better understand the role of obesity in HF.

Therefore, the main objective of this study was to identify and validate the differentially expressed miRNAs involved in the obesity paradox and to evaluate the mechanisms potentially modulated by their action.

## Materials and methods

### Patients and controls

Patients were recruited at the Heart Failure and Transplant Clinic of our institution from March 2012 to December 2013 for a prospective cohort from our group. Previous data from this study had been published [10], and samples were stored in a -80°C freezer for future analyses. Subjects with stable HF, 18 to 80 years old, and a left ventricular ejection fraction (LVEF) lower than 45% were included. Exclusion criteria for HF patients were as follows: (I) episode of decompensation within the previous 30 days, (II) acute coronary syndrome within the previous three months, and (III) presence of implantable devices such as pacemakers, implantable cardioverter defibrillators, and cardiac resynchronization therapy-defibrillators (CRT-D), which were not eligible for bioelectrical impedance analysis.

For obese patients, the inclusion criteria were BMI (body weight in kg/ height in meters squared) ≥30 kg/m^2^ and also the World Health Organization criteria for obesity based on percent body fat (> 25% in men and > 35% in women) [2], as measured by bioelectrical impedance analyzer (Model 450, Tetrapolar, Biodynamics). Lean subjects were included if BMI was < 25 kg/m^2^ and body fat was ≤22% in men and ≤32% in women [11].

Healthy controls were selected from donors of the Blood Center of our institution and also selected at the Echocardiography Laboratory, from patients showing a normal exam. All controls had no history of structural cardiac disease or symptoms of HF, and body composition was classified as normal. Exclusion criteria for both HF patients and controls included cell dyscrasias, active inflammation, malignant disease, and severe hepatic or renal disease (creatinine > 3 mg/dL). Subjects were divided into three groups: (I) lean healthy controls, (II) lean with HF, and (III) obese with HF. All groups were matched for age and sex, and groups II and III were also matched for LVEF, HF etiology (ischemic and non-ischemic), New York Heart Association (NYHA) class, and use of beta-blockers, and angiotensin-converting enzyme (ACE) inhibitor or angiotensin receptor blocker (ARB).

### Data collection

Demographic data, clinical history, comorbidities, echocardiographic, electrocardiographic, and laboratory data were collected. Body mass index was calculated as weight in kilograms divided by squared height in meters (kg/m^2^). Bioimpedance analysis was performed with tetrapolar bioimpedance of Biodynamics, model 450. Waist circumference was measured by a trained examiner and determined on the lesser curvature located between the ribs and the iliac crest after the expiration of the patient. Data were collected retrospectively and assessed in January of 2020. The author matched samples to dataset information by participant ID number.

### Sample preparation

Peripheral venous blood samples from all subjects were collected in EDTA-coated tubes. Blood samples were centrifuged at 1,500 rpm for 15 minutes at 4°C within one hour from the collection and stored in 500 μL microtubes at -80°C for posterior analysis.

### Microarray analysis

For microarray, total RNA was extracted from 396 μL of plasma spiked-in with 4 μL of cel-miR-39-3p. Then, 1.2 mL of Trizol LS reagent was added and incubated for 5 minutes. Next, 0.32 mL of chloroform was added, incubated for 15 minutes, and centrifuged at 12,000 x g at 4°C for 15 minutes. Then, we removed the supernatant, added 0.8 mL of isopropyl alcohol, incubated for 10 minutes, and centrifuged at 12,000 x g at 4°C for 10 minutes. After removing the isopropyl alcohol, 1.6 mL of 75% ethanol was added and centrifuged at 7,500 x g, at 4°C, for 5 minutes. Dry for 10 minutes and resuspend in 16 uL of RNase-free water, placed in a dry bath at 58°C for 10 min. The concentration of miRNAs was determined by spectrophotometric analysis (NanoDrop 1000, Thermo Scientific, Wilmington, DE, USA).

Affymetrix GeneChip miRNA 4.0 Array was used to scan the miRNAs in patient samples. RNA labeling and array hybridization was performed following the manufacturer’s instruction. Differential analysis of the miRNAs was performed using Affymetrix GeneChip miRNA array technology version 4.0 (Thermo Fisher Scientific, Santa Clara, CA, USA), according to the manufacturer’s specifications on total RNA extracted from plasma. The array contains probe sets for 2,578 mature human miRNAs. Total RNA (160 ng), including miRNAs, was biotin-labeled using the FlashTagTM Biotin HSR RNA Labeling kit (Affymetrix, Genisphere, Hatfield, PA, USA) and the samples were then hybridized overnight using the GeneChip Hybridization Oven 640 (Affymetrix, Santa Clara, CA, USA) at 48°C. The arrays were then washed and stained in the GeneChip Fluidics Station 450 (Affymetrix, Santa Clara, CA, USA). The arrays were scanned using a GeneChip Scanner 3000 7G (Affymetrix, Santa Clara, CA, USA) and the signal values were evaluated using the Expression Console Software (EC) v1.2 (Affymetrix by Thermo Fisher Scientific). Intensity values (presence/absence values) and signal histograms of each hybridization were quality-checked.

Raw data were pre-processed using the Robust Multiarray Average (RMA) method, which applies background correction, log2 transformation, and quantile normalization. The batch effect was adjusted with Surrogate Variable Analysis through the sva R package [12]. Pairwise differential expression analysis was carried out with the Limma R package [13], and a p-value < 0.01 was adopted to identify differentially expressed miRNAs.

The data used in this publication have been deposited in NCBI’s Gene Expression Omnibus and are accessible through GEO Series accession number GSE288767 (https://www.ncbi.nlm.nih.gov/geo/query/acc.cgi?acc=GSE288767).

### RT-qPCR

Circulating miRNA from 198 μL of plasma spiked-in with 2 μL of cel-miR-39-3p was extracted using the miRNeasy Serum/Plasma Kit. We added 1 mL of QIAzol and incubated for 5 minutes. Then, 200 μL of chloroform were added, vortexed for 15 seconds, incubated for 3 minutes, and centrifuged for 15 minutes at 12,000 x g at 4°C. We removed the supernatant and added 1.5 volumes of 100% ethanol. Afterward, we pipetted 700 μL into the extraction column and centrifuged at 8,000 x g for 60 seconds at room temperature (15–25°C). We then added 700 μL of RWT buffer to the column and centrifuged for 60 s at 8,000 x g. Next, we pipetted 500 μL of RPE buffer onto the column and centrifuged for 60 s at 8,000 x g. Pipette 500 μL of 80% ethanol onto the column and then centrifuged for 2 minutes at 8,000 x g. Afterward, we centrifuged the column in a new tube for another 5 minutes to eliminate the rest of the ethanol. In the end, we added 14 μL of RNase-free water and centrifuged at 8,000 x g for 60 s at room temperature (15–25°C). The concentration of miRNAs was determined by spectrophotometric analysis (NanoDrop 1000).

RT-PCR was performed using the StepOnePlus™ equipment. For the poly-A tail reaction, 1.75 μL of sample in 2.75 μL of Poly(A) Reaction Mix was added to each tube. Then, the samples were placed into a thermocycler for polyadenylation at a temperature of 37°C for 45 minutes and stopped the reaction at 65°C for 10 minutes. Next, the adapter ligation reaction was performed with 7.5 μL of Ligation Reaction Mix, at a temperature of 16°C for 60 minutes. After, reverse transcription (RT) was performed using 11.25 μL of RT Reaction Mix at 42°C for 15 minutes and stop reaction at 85°C for 5 minutes. The miR-Amp reaction was performed with 3.75 μL of the RT product in 33.75 μL of the miR-Amp Reaction Mix, activating the enzyme at 95°C for 5 minutes, denaturing at 95°C for 3 seconds, annealing, and extension at 60°C for 30 seconds (14 cycles - denaturation, annealing, and extension phase), and stop of the reaction at 99°C for 10 minutes.

Each amplification reaction was prepared with 5 μL of TaqMan® Fast Advanced Master Mix (2X), 0.5 μL TaqMan® Advanced miRNA Assay (20X), and 2 μL of RNase-free water, in MicroAmp® Flat Optical 48-Well Reaction Plate (0.1 mL); enzyme activation at 95°C, for 20 seconds, denaturation at 95°C for 1 second; annealing and extension at 60°C for 20 seconds (40 cycles - denaturation, annealing, and extension phase).

In brief, data from qRT-PCR were normalized to the reference control miRNA, cel-miR-39-3p, using the ΔCT method. The relative expression levels for each individual miRNA were calculated using the following mathematical formula: ΔCT = CTsample – CTcel-miR-39-3p. The ΔΔCT for each miRNA was calculated using the formula: ΔΔCT = ΔCTsample – ΔCT mean of control group. These values were transformed into quantities using the formula 2^−ΔΔCT and are presented as fold-change relative to the internal control.

### Target and pathway analysis of validated miRNAs

The differentially expressed miRNAs validated through qRT-PCR, hsa-miR-451a, hsa-miR-22-3p, and hsa-miR-548ac were submitted to target analysis using the miRTarBase release 8.0 [16] and TarBase v8.0 [14]. Both databases contain experimentally validated miRNA-target interactions and were explored applying the following filtering criteria: (I) for miRTarBase, we restricted for interactions classified as functional (both weak and strong); (II) for TarBase, we considered only interactions classified as “positive” and related to a direct association between miRNA and their target gene. Moreover, for TarBase, all interactions supported by HITS-CLIP and PAR-CLIP high-throughput experiments were further analyzed to count the number of supporting experiments per interaction. To mitigate the impact of publication bias (i.e., the fact that some miRNAs are studied more frequently than others), we applied a uniform selection criteria for all analyzed miRNA-target interactions. Specifically, only interactions with at least 2 supporting pieces of evidence were retained, regardless of the miRNA. This approach was applied to hsa-miR-451a, hsa-miR-22-3p, and hsa-miR-548ac, ensuring consistency in the analysis process.

The union of miRNA-target interactions retrieved from both databases was used as the interactome of the validated miRNAs. The functional enrichment was quantitatively assessed (p-value) using a hypergeometric distribution. Multiple test correction was also implemented by applying the False Discovery Rate (FDR) algorithm [17] at a significance level of p<0.05. We used the CluePedia: A ClueGO plugin for pathway insights using integrated experimental and *in silico* data v.2.5.7 [18,19], with the terms or pathways consulted in the KEGG database [15].

### Ethical considerations

The study protocol was conducted according to the principles outlined in the Declaration of Helsinki and was approved by our institution’s Ethics Committee in Research under the numbers 120084 and 19470 with certification of ethical evaluation number CAAE: 2414481.1.0000.5327. We obtained written informed consent from all patients prior to their inclusion in the study.

### Statistical analysis

Quantitative data were described as mean and standard deviation and frequencies as absolute values and percentages. In the RT-PCR analysis, the differential expression of miRNAs was analyzed using the geometric mean and standard deviation. For group comparisons, we used the Kruskal-Wallis test and Dunn’s multiple comparison test, with a p-value < 0.05.

## Results

### Discovery cohort

#### Baseline characteristics

A total of 20 HF patients (10 obese and 10 lean) and 10 healthy controls were enrolled for the discovery phase. Clinical and laboratory characteristics of patients and controls are shown in Table 1. The three groups were balanced for age, sex, and ethnicity. The obese-HF group presented the adiposity parameters characteristic of the population with increased body weight, BMI, fat percentage, and abdominal circumference. Both HF groups had severe systolic dysfunction and mild functional limitation. Most patients were under a standard therapeutic regimen with beta-blockers and angiotensin-converting enzyme inhibitors.

**Table 1.**
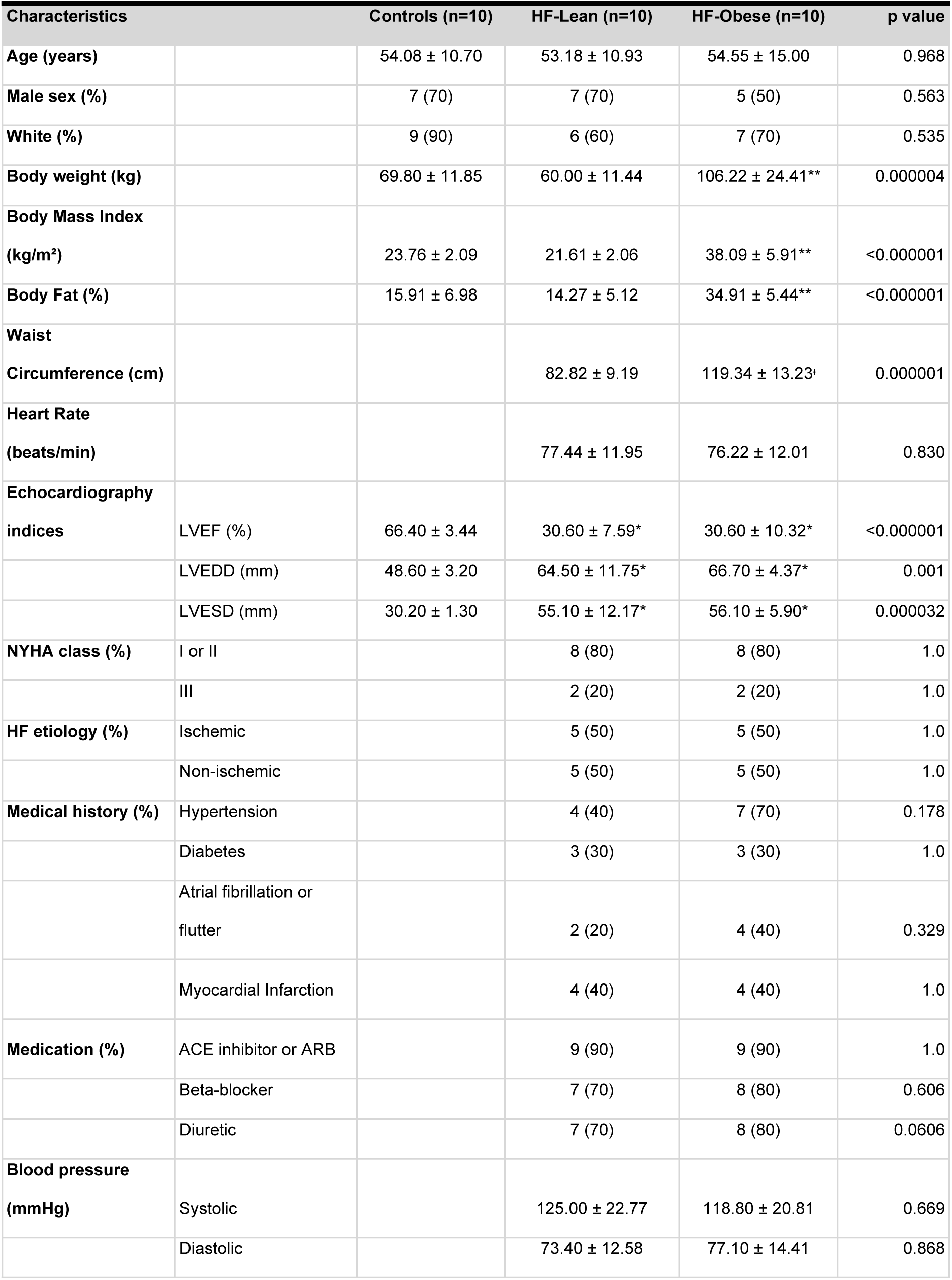
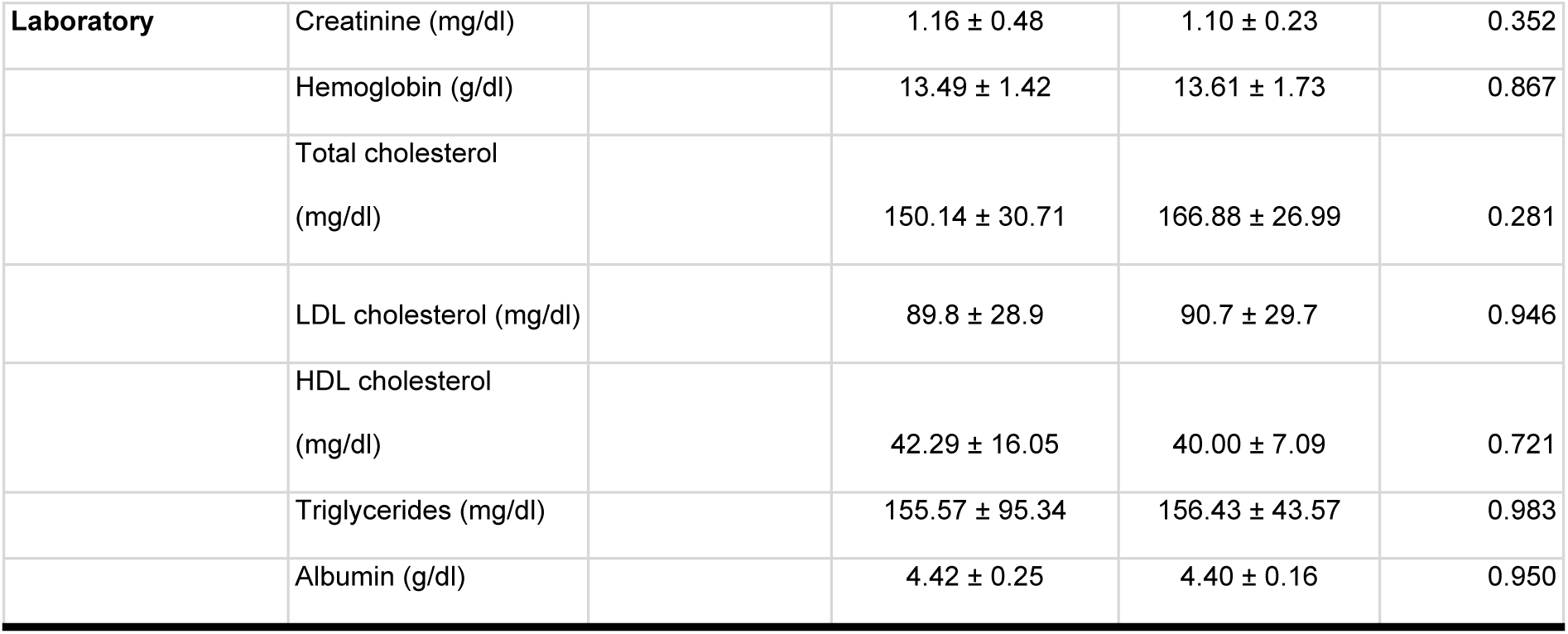
Discovery cohort baseline characteristics. Table 1 presents the baseline characteristics of the discovery cohort, which includes three groups: Control (n=10), Heart Failure Lean (HF-Lean, n=10), and Heart Failure Obese (HF-Obese, n=10). The p-values indicate statistically significant differences between the groups, with values less than 0.05 (<0.05). *Statistically significant difference between the HF groups vs. Control. **Statistically significant difference between the Obese group vs. Lean.

#### Microarray results

In the microarray analysis of 2,578 human miRNAs, 48 miRNAs were found to be differentially expressed in the comparison between HF-lean vs controls, 21 miRNAs in the comparison between HF-obese vs healthy controls, and 6 miRNAs differentially expressed in the comparison between HF-obese vs HF-lean. Afterwards, we built a Venn diagram to evaluate the overlaps of each comparison. Eleven HF-related miRNAs and one obesity-related miRNA were observed. Among the HF-related miRNAs we observed up-regulated miR-378h, miR-22-3p, miR-140-3p, miR-106b-5p, miR-3196, and miR-6125 and down-regulated miR-548ac, miR-8084, miR-3128, miR-574-5p, and miR-3201. Obesity-related miR-451a was up-regulated (Fig 1).

**Fig 1.**
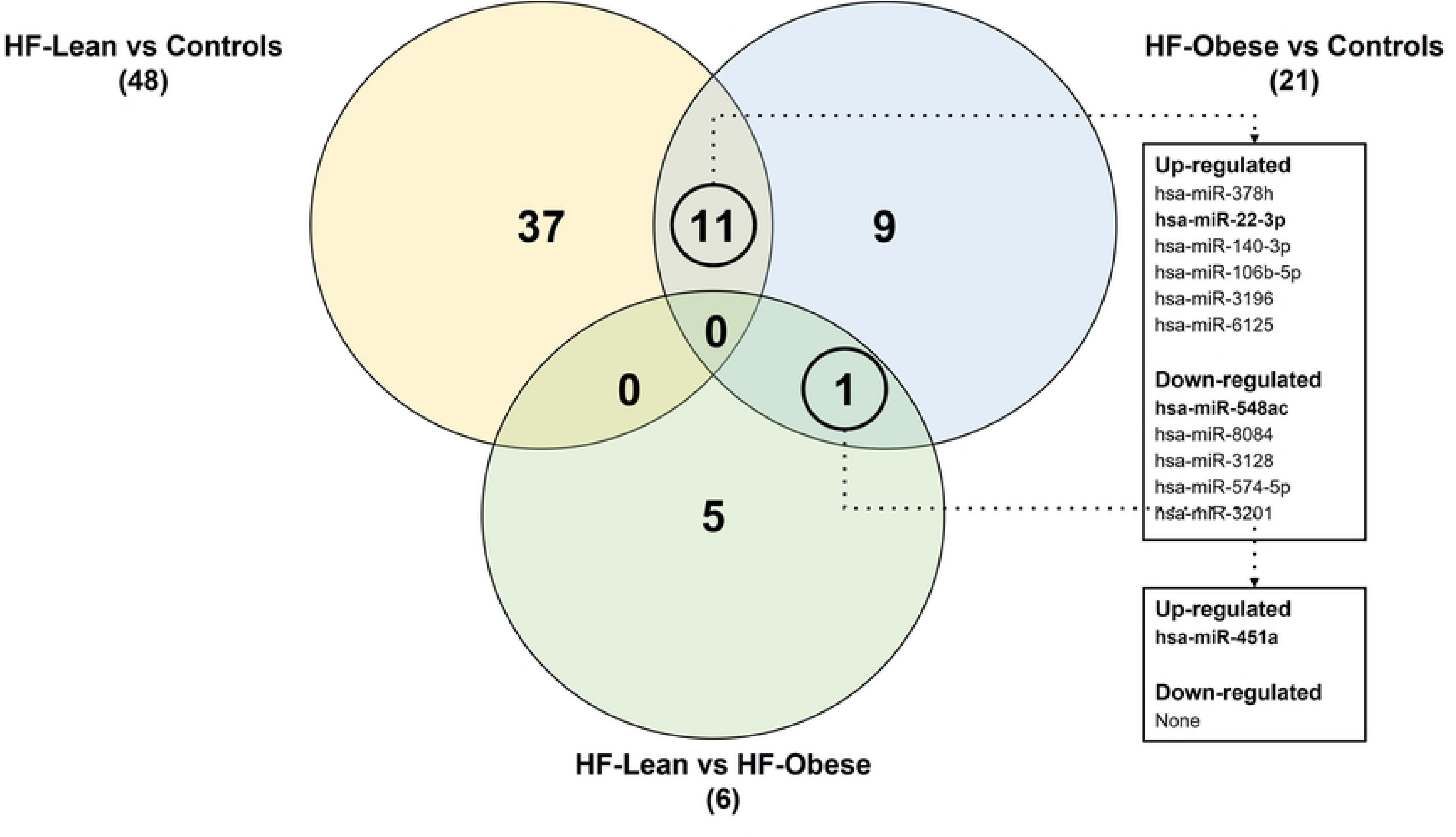
Venn Diagram of differentially expressed miRNAs. Venn diagram illustrating differentially expressed microRNAs across three comparisons: HF-Lean vs Controls, HF-Obese vs Controls, and HF-Lean vs HF-Obese. The numbers represent unique and shared differentially expressed microRNAs. HF-Lean vs Controls had 48, of which 37 were unique. HF-Obese vs Controls showed 21, with 9 unique. HF-Lean vs HF-Obese identified 6, with 5 unique. Eleven microRNAs were commonly differentially expressed in HF-Lean and HF-Obese compared to controls, while only one was shared between HF-Obese vs Control and HF-Lean vs HF-Obese. The adjacent table lists up- and down-regulated microRNAs for each comparison.

### Validation cohort

#### Baseline characteristics

A total of 80 subjects, 61 HF patients (26 obese and 35 lean), and 19 healthy controls were enrolled for the validation phase of the study. Clinical and laboratory characteristics of patients and controls are shown in Table 2. The obese HF group was well characterized for obesity parameters. Both HF groups had severe systolic dysfunction and mild functional limitation. Most patients were under a standard therapeutic regimen with beta-blockers and angiotensin-converting enzyme inhibitors. There were no significant differences in clinical variables between both HF groups except for hypertension.

**Table 2.**
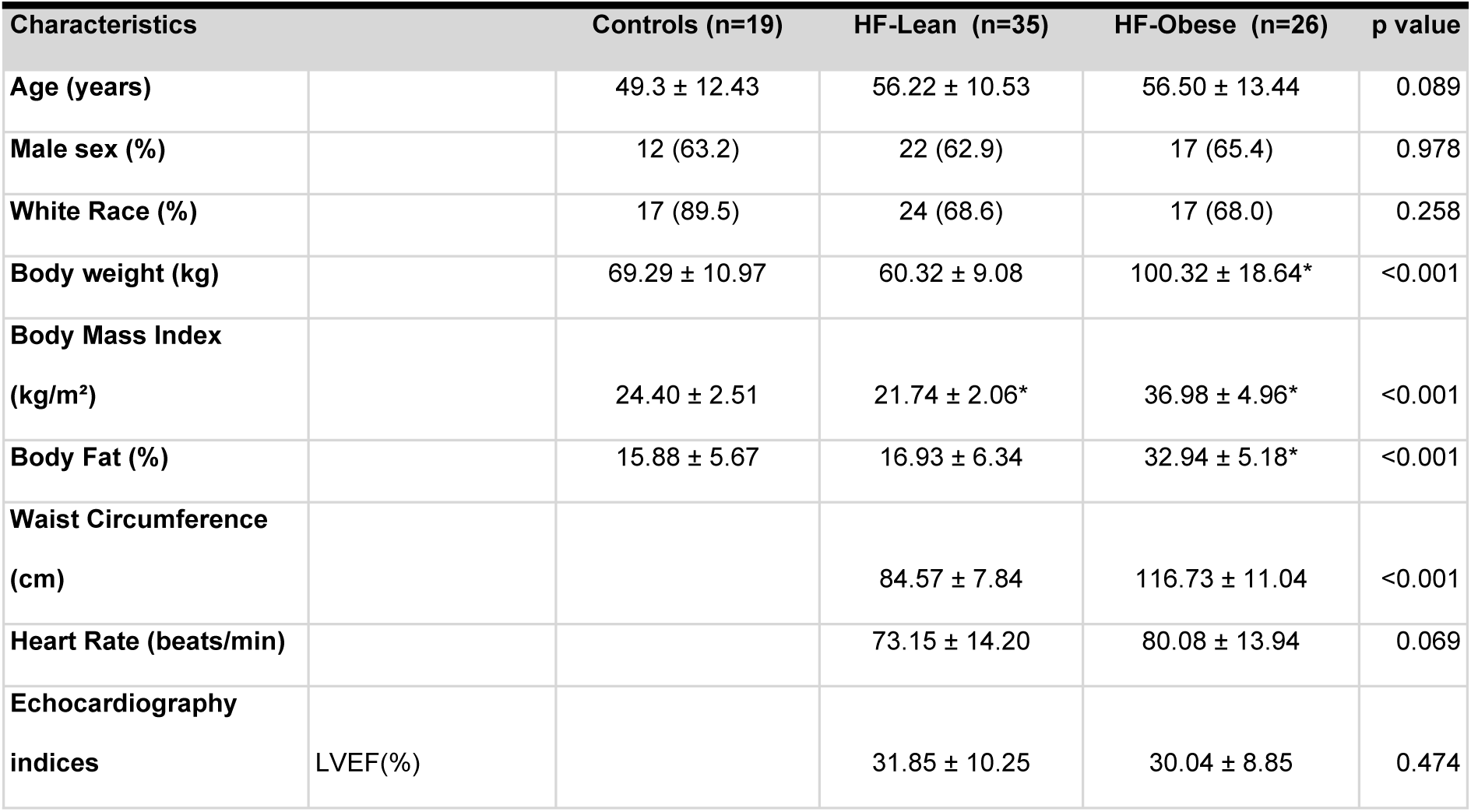
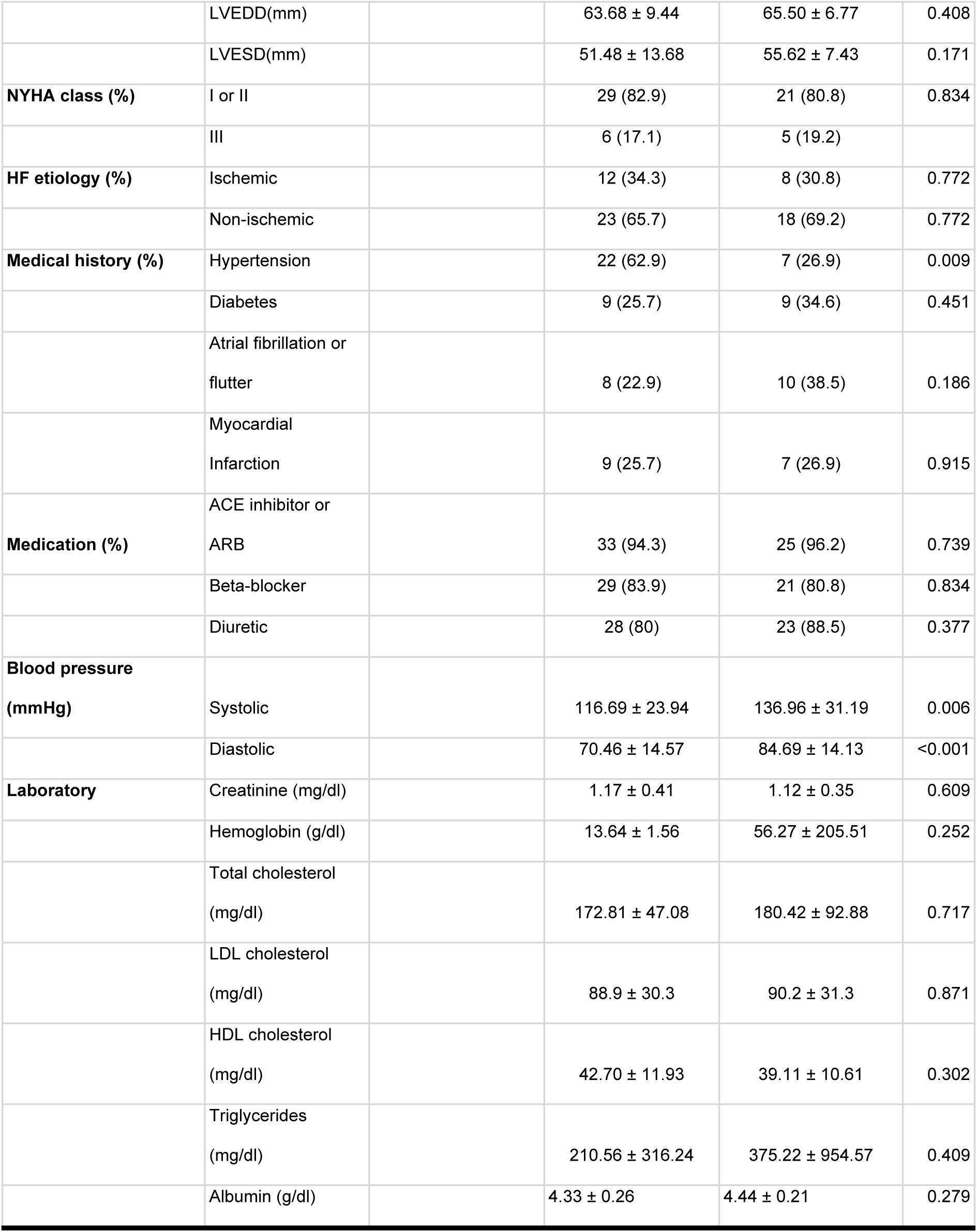
Validation cohort baseline characteristics. Table 2: Baseline characteristics of the validation cohort. This table presents the baseline characteristics of the three groups in the validation cohort: Controls (n=19), HF-Lean (n=35), and HF-Obese (n=26). Values are presented as mean ± standard deviation. The p-value column indicates statistically significant differences between the groups. * Difference between HF groups vs. Control, p-value < 0.05.

#### Validation results

Based on microarray results, we ranked miRNAs by effect size (-1; 1 fold-change), lowest p-value (<0.01), and biological plausibility to prioritize candidates for validation. Subsequently, miRNAs -451a, -22-3p, and -548ac were selected for further analysis. Notably, both miR-451a and miR-22-3p were up-regulated in both HF groups compared to the control, validated in our sample. However, miR-548ac was down-regulated in the microarray and up-regulated in the validation sample, not validating the initial result (Fig 2).

**Fig 2.**
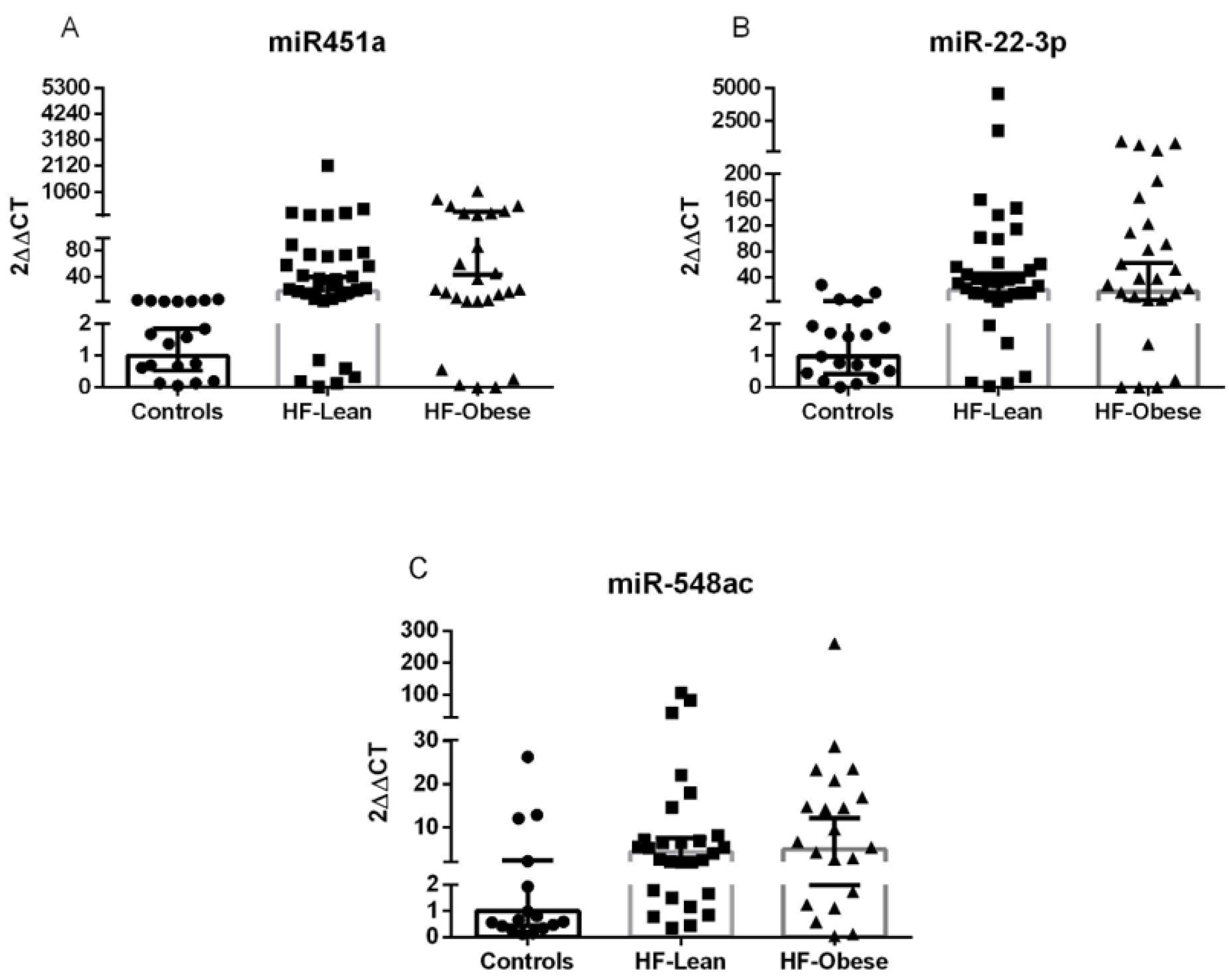
miRNAS validated by qRT-PCR. Validation of selected microRNA expression in experimental groups. The relative expression of microRNAs miR-451a (A), miR-22-3p (B), and miR-548ac (C) was analyzed in the Control, HF-Lean, and HF-Obese groups. The microRNAs were selected based on microarray results. miR-451a and miR-22-3p were upregulated in the HF groups, validating the microarray, while miR-548ac was not confirmed.

### Construction of HF-miR-gene network

After identifying and ranking the differentially expressed miRNAs, we performed an interaction analysis among miRNAs-genes-pathways. The interaction of these miRNAs resulted in a network with 937 genes involved. We then filtered genes with single interaction with miRNA, which yielded a network with 36 genes (Fig 3). We selected the signaling pathways that interacted with these genes and filtered them by FDR < 0.05, totaling 9 pathways (S1 Table). Finally, we assessed the most frequent genes across these pathways and ranked them. The most frequent genes were AKT1, MAPK1, GRB2, IGF1R, PTEN, ESR1, HSPA1B, MAP3K1, and ZFHX3. These genes are associated with different biological processes, such as apoptosis, carbohydrate metabolism, neurogenesis, sugar transport, translation regulation, transport, cell cycle, actin cytoskeleton reorganization, cardiac neural crest cell development involved in heart development, and aging.

**Fig 3.**
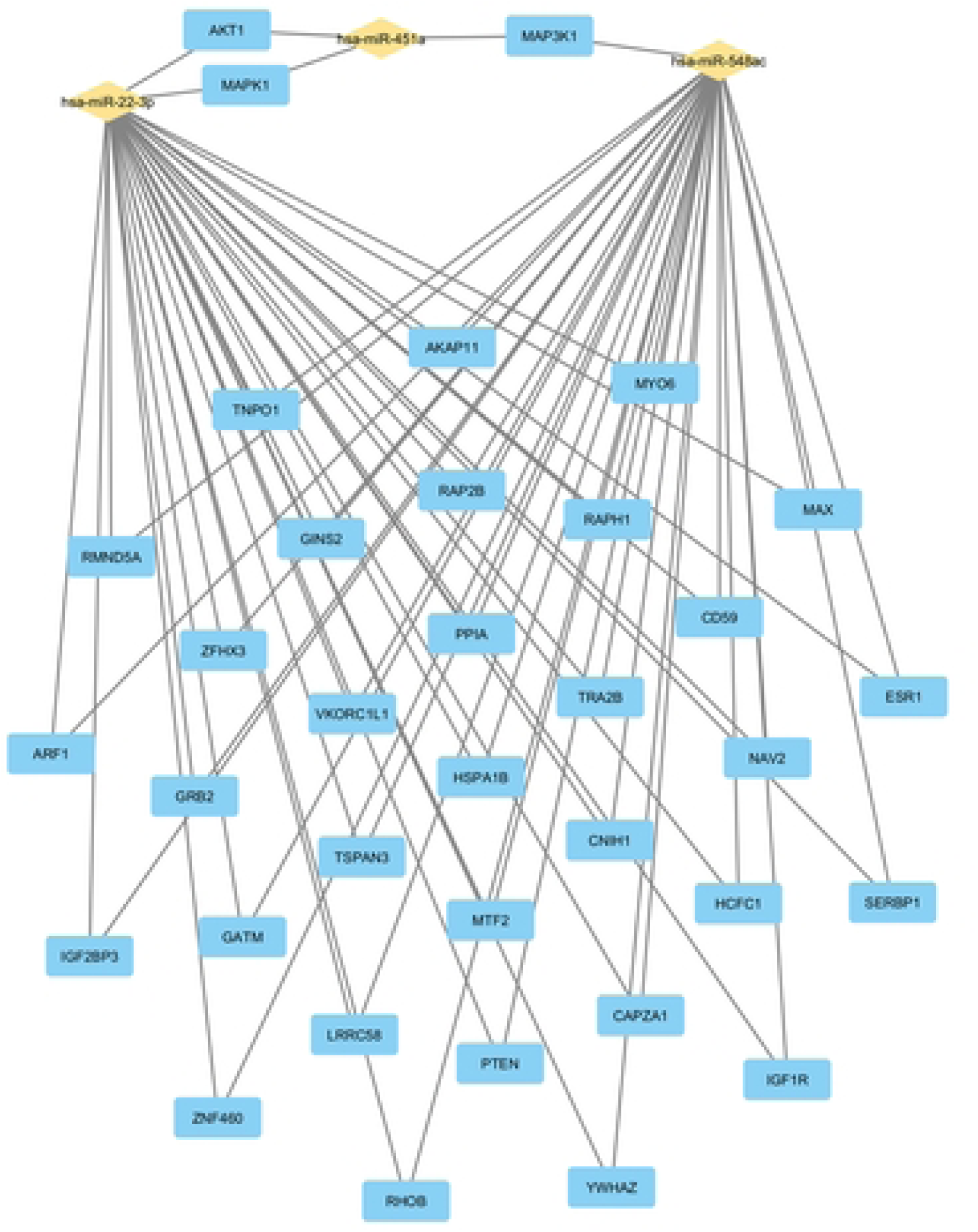
miRNAs-gene network. Interaction network between microRNAs and target genes. The figure shows three microRNAs — hsa-miR-22-3p, hsa-miR-451a, and hsa-miR-548ac (in yellow) — and their respective target genes (in blue). The lines connecting the miRNAs to the genes represent regulatory interactions, indicating that these miRNAs control the expression of the linked genes.

## Discussion

In the discovery phase of the present study, of the 2,578 human miRNAs, 48 were differentially expressed - 11 related to HF and one to obesity. After evaluating the effect size, p-value, and biological plausibility of the differentially expressed miRNAs, we chose three of them for validation - miR-451a, miR-22-3p, and miR-548ac. In the validation phase, miRNAs -451a and -22-3p were up-regulated in the HF groups, regardless of obesity. The miR-548ac showed to be divergent between the discovery (down-regulated) and validation (up-regulated) phases in the HF groups compared to the control group. Through *in silico* analysis, we observed that this group of miRNAs is associated with different important biological processes that participate in metabolic, morphological, and functional activities in HF through genes such as *AKT1*, *MAPK1*, *GRB2*, and *IGF1R*.

Our results show that miR-451a is up-regulated in the HF groups regardless of obesity compared to the control group. Interaction analysis showed that miR-451a is associated with *AKT1*, *MAPK1*, and *MAP3K1* genes. These genes interact with biological processes, such as apoptosis, metabolism, proliferation, cell survival, growth, and angiogenesis, which are fundamental for cardiac remodeling and function. As in our population, patients with hypertrophic cardiomyopathy have increased levels of miR-451a [20], showing an association with cardiac remodeling. Some preclinical studies have shown that the increased expression of miR-451a prevents activation of matrix metalloproteinases 2 and 9 in human cardiomyocytes during pathological stress stimulation [21] and attenuates cardiac fibrosis and angiotensin II-induced inflammation in mice [22]. It is possible that miR-451a has a cardioprotective role in the heart disease scenario.

Our analyses showed that the miR-22-3p was up-regulated in both HF groups compared to the control group. Based on interaction analysis, we found that miR-22-3p is associated with *AKT1* and *MAPK1* genes, in conjunction with miR-451a, suggesting a synergy of these miRNAs in gene expression regulation, and consequently, modulating the same biological processes. In the Bio-SHiFT study [23], it was observed that miR-22-3p is an independent marker, inversely associated with primary outcomes such as heart failure, hospitalization, cardiovascular mortality, cardiac transplantation, and LVAD implantation. A recent preclinical study [24] showed that upregulation of miR-22-3p generated inhibitory effects on cell proliferation and collagen deposition in cardiac fibroblasts treated with Ang II. This evidence suggests that miR-22-3p is associated with cardioprotective factors that should be further explored.

The results of miR-548ac were divergent between the discovery and validation phases. While this miRNA was down-regulated in our microarray analysis, we observed its up-regulation in the RT-qPCR analysis, showing an increased expression in patients with HF compared to the control group. The miRNA target analysis highlighted genes *MAPK1*, *GRB2*, and *IGF1R* as potential targets for this miRNA. These genes participate in the activation of biological processes such as cell proliferation, cell surface growth factor receptors, the Ras signaling pathway, cell growth, and survival control. To date, our study appears to be the first to analyze miR-548ac in patients with HF. Previously, only studies evaluating its expression in the context of rheumatic diseases [25] and cancer [26–28] were reported. Our data suggest that exploring miR-548ac and the mechanisms involved in patients with HF may be relevant.

The main limitation of our study is that we did not include a group of healthy obese individuals in our analyses.

## Conclusion

Three circulating miRNA were identified using microarray technology, miRNAs -451a, -22-3p, and -548ac. The validation of the differential expression was confirmed for miRNAs miR-451a and miR-22-3p, with both showing up-regulation in HF regardless of obesity. No miRNAs were associated with either HF and obesity. These miRNAs are associated with the *AKT1* and *MAPK1* genes, which participate in biological processes including metabolism, proliferation, cell survival, growth, and angiogenesis, important in HF. Further studies are warranted to clarify the interaction of these miRNA pathways into specific HF pathogenic mechanisms.

## Acknowledgments (Grants)

This study was supported by Conselho Nacional de Desenvolvimento Científico e Tecnológico (CNPq) through postgraduate students scholarship and by Coordenação de Aperfeiçoamento de Pessoal de Nível Superior – Brasil (CAPES, Coordination of Superior Level Staff Improvement) – Finance Code 001. We also acknowledge the partial support from Fundação de Amparo à Pesquisa do Estado do Rio Grande do Sul (FAPERGS) through grant nr. 16/2551-0000520-6.

## References

1. Prospective Studies Collaboration, Whitlock G, Lewington S, Sherliker P, Clarke R, Emberson J, et al. Body-mass index and cause-specific mortality in 900 000 adults: collaborative analyses of 57 prospective studies. Lancet. 2009;373: 1083–1096.

2. Cornier M-A, Després J-P, Davis N, Grossniklaus DA, Klein S, Lamarche B, et al. Assessing adiposity: a scientific statement from the American Heart Association. Circulation. 2011;124: 1996–2019.

3. Kenchaiah S, Sesso HD, Gaziano JM. Body mass index and vigorous physical activity and the risk of heart failure among men. Circulation. 2009;119: 44–52.

4. Joyce E, Lala A, Stevens SR, Cooper LB, AbouEzzeddine OF, Groarke JD, et al. Prevalence, Profile, and Prognosis of Severe Obesity in Contemporary Hospitalized Heart Failure Trial Populations. JACC Heart Fail. 2016;4: 923–931.

5. Oktay AA, Lavie CJ, Kokkinos PF, Parto P, Pandey A, Ventura HO. The Interaction of Cardiorespiratory Fitness With Obesity and the Obesity Paradox in Cardiovascular Disease. Prog Cardiovasc Dis. 2017;60: 30–44.

6. Padwal R, McAlister FA, McMurray JJV, Cowie MR, Rich M, Pocock S, et al. The obesity paradox in heart failure patients with preserved versus reduced ejection fraction: a meta-analysis of individual patient data. Int J Obes. 2014;38: 1110– 1114.

7. Sharma A, Lavie CJ, Borer JS, Vallakati A, Goel S, Lopez-Jimenez F, et al. Meta-analysis of the relation of body mass index to all-cause and cardiovascular mortality and hospitalization in patients with chronic heart failure. Am J Cardiol. 2015;115: 1428–1434.

8. Thaker VV. GENETIC AND EPIGENETIC CAUSES OF OBESITY. Adolesc Med State Art Rev. 2017;28: 379–405.

9. Kim SY, Morales CR, Gillette TG, Hill JA. Epigenetic regulation in heart failure. Curr Opin Cardiol. 2016;31: 255–265.

10. Thomé JG, Mendoza MR, Cheuiche AV, La Porta VL, Silvello D, Dos Santos KG, et al. Circulating microRNAs in obese and lean heart failure patients: A case-control study with computational target prediction analysis. Gene. 2015;574: 1– 10.

11. Pi-Sunyer FX. Obesity: criteria and classification. Proc Nutr Soc. 2000;59: 505– 509.

12. Leek JT, Johnson WE, Parker HS, Jaffe AE, Storey JD. The sva package for removing batch effects and other unwanted variation in high-throughput experiments. Bioinformatics. 2012;28: 882–883.

13. Ritchie ME, Phipson B, Wu D, Hu Y, Law CW, Shi W, et al. limma powers differential expression analyses for RNA-sequencing and microarray studies. Nucleic Acids Res. 2015;43: e47.

14. Karagkouni D, Paraskevopoulou MD, Chatzopoulos S, Vlachos IS, Tastsoglou S, Kanellos I, et al. DIANA-TarBase v8: a decade-long collection of experimentally supported miRNA-gene interactions. Nucleic Acids Res. 2018;46: D239–D245.

15. Kanehisa M, Goto S, Kawashima S, Nakaya A. The KEGG databases at GenomeNet. Nucleic Acids Res. 2002;30: 42–46.

16. Huang H-Y, Lin Y-C-D, Li J, Huang K-Y, Shrestha S, Hong H-C, et al. miRTarBase 2020: updates to the experimentally validated microRNA-target interaction database. Nucleic Acids Res. 2020;48: D148–D154.

17. Benjamini Y, Hochberg Y. Controlling the false discovery rate: A practical and powerful approach to multiple testing. J R Stat Soc. 1995;57: 289–300.

18. Bindea G, Mlecnik B, Hackl H, Charoentong P, Tosolini M, Kirilovsky A, et al. ClueGO: a Cytoscape plug-in to decipher functionally grouped gene ontology and pathway annotation networks. Bioinformatics. 2009;25: 1091–1093.

19. Bindea G, Galon J, Mlecnik B. CluePedia Cytoscape plugin: pathway insights using integrated experimental and in silico data. Bioinformatics. 2013;29: 661– 663.

20. Sonsöz MR, Yilmaz M, Cevik E, Orta H, Bilge AK, Elitok A, et al. Circulating Levels of MicroRNAs in Hypertrophic Cardiomyopathy: The Relationship With Left Ventricular Hypertrophy, Left Atrial Dilatation and Ventricular Depolarisation-Repolarisation Parameters. Heart Lung Circ. 2022;31: 199–206.

21. Scrimgeour NR, Wrobel A, Pinho MJ, Høydal MA. microRNA-451a prevents activation of matrix metalloproteinases 2 and 9 in human cardiomyocytes during pathological stress stimulation. Am J Physiol Cell Physiol. 2020;318: C94–C102.

22. Deng H-Y, He Z-Y, Dong Z-C, Zhang Y-L, Han X, Li H-H. MicroRNA-451a attenuates angiotensin II-induced cardiac fibrosis and inflammation by directly targeting T-box1. J Physiol Biochem. 2022;78: 257–269.

23. van Boven N, Akkerhuis KM, Anroedh SS, Rizopoulos D, Pinto Y, Battes LC, et al. Serially measured circulating miR-22-3p is a biomarker for adverse clinical outcome in patients with chronic heart failure: The Bio-SHiFT study. Int J Cardiol. 2017;235: 124–132.

24. Zhao X-S, Ren Y, Wu Y, Ren H-K, Chen H. MiR-30b-5p and miR-22-3p restrain the fibrogenesis of post-myocardial infarction in mice via targeting PTAFR. Eur Rev Med Pharmacol Sci. 2020;24: 3993–4004.

25. Guo S, Jin Y, Zhou J, Zhu Q, Jiang T, Bian Y, et al. MicroRNA Variants and HLA-miRNA Interactions are Novel Rheumatoid Arthritis Susceptibility Factors. Front Genet. 2021;12: 747274.

26. Song F, Yang Y, Liu J. MicroRNA-548ac induces apoptosis in laryngeal squamous cell carcinoma cells by targeting transmembrane protein 158. Oncol Lett. 2020;20: 69.

27. Lyu P, Hao Z, Zhang H, Li J. Identifying pancreatic cancer-associated miRNAs using weighted gene co-expression network analysis. Oncol Lett. 2022;24: 297.

28. Barquet-Muñoz SA, Pedroza-Torres A, Perez-Plasencia C, Montaño S, Gallardo-Alvarado L, Pérez-Montiel D, et al. microRNA Profile Associated with Positive Lymph Node Metastasis in Early-Stage Cervical Cancer. Curr Oncol. 2022;29: 243–254.

